# Females Adapt to Dietary Protein Restriction on Enhanced Gut-Brain Axis during Aging

**DOI:** 10.64898/2026.06.29.735363

**Authors:** Prerana Vaddi, Jose G. Lugo, Khristina E. Young, Yvann Batamack, Michael Donkor, Ahivital Artison, Amy Christensen, Christian J. Pike, Cristal M. Hill

## Abstract

Growing evidence supports a critical role for the gut-brain axis in regulating metabolic health, inflammation, and cognitive function during aging. Age-associated gut dysbiosis has been linked to metabolic dysfunction and cognitive decline, with females exhibiting increased susceptibility to these age-related impairments. Diet is a major determinant of gut microbiome composition and function. Previous studies from our laboratory demonstrated that dietary protein restriction (DPR) induces fibroblast growth factor 21 (FGF21), improves metabolic health, and extends lifespan in male mice. However, the effects of DPR on the gut microbiome and associated health outcomes in aged female mice remain poorly understood. Female mice were assigned at 16 months of age to either a normal-protein (NP) or low-protein (LP) diet for 26 weeks. Metabolic assessments included food intake, fasting glucose concentrations, and glucose tolerance testing. Senescence-associated markers in mesenteric white adipose tissue (mWAT), fecal microbiome composition, and behavioral outcomes were evaluated to determine relationships among dietary protein intake, microbial communities, metabolic health, and cognitive function. Low-protein diet significantly improved metabolic health in aged female mice, as evidenced by improved glucose regulation. Microbiome analyses revealed increased abundance of *Akkermansia* at 17 months and *Faecalibaculum* in LP-fed animals at 22 months of age. More so, functional profiling and gene set enrichment analyses indicated enrichment of microbial pathways associated with membrane integrity and metal ion binding. Lastly, LP-fed female mice displayed improved memory performance at 22 months of age compared with age-matched NP-fed controls. Collectively, these findings demonstrate that DPR remodels the gut microbiome and improves metabolic and cognitive health in aged female mice. The observed microbial adaptations may contribute to the beneficial effects of DPR on aging-related physiology, highlighting the gut microbiome as a potential mediator of dietary interventions that promote healthy aging.

## Introduction

Aging is associated with significant changes that negatively impact the gut microbiota, leading to a condition known as gut dysbiosis(^1–3^). This imbalance in microbial composition can promote inflammation, impair metabolism, and decline cognition, all of which contribute to age-related diseases. Over the past 10 years, the gut has gained attention as a central organ connecting metabolic health and learning behavior in response to dietary restriction paradigms such as calorie restriction, fasting, or altering amino acid intake(^4–6^). Similarly, among these paradigms, dietary protein restriction (DPR), has emerged as a promising strategy to benefit metabolic health in invertebrates and vertebrates, as well as extend lifespan in rodents(^7–11^).

As the primary site of nutrient absorption and a critical interface between host and environment, the gut microbiome plays an essential role in shaping general health, including cognition. The gut-brain axis, an intricate communication network between the gastrointestinal tract and the central nervous system, modulates brain health through neural (e.g., vagal), immune, and endocrine pathways. Notably, microbial metabolites like short-chain fatty acids, trimethylamine N-oxide, polyunsaturated fatty acids, tryptophan, and indole derivatives can influence microglial function, blood-brain barrier integrity, and synaptic transmission(^12–14^). Studies have shown that DPR reduces tau accumulation in Alzheimer’s disease-like murine models in females. Though these studies show that DPR can improve specific AD markers, the mechanisms through which DPR definitively influences brain health in the absence of neurodegeneration are still unclear.

Prior studies investigating the relationship between the gut microbiome and DPR have shown that DPR remodels the microbiome in six-month-old male mice, resulting in unique changes in microbial composition and corresponding improvements in metabolic health and hepatic adaption(^15^). Additionally, a recent study examining fiber supplementation alongside DPR in two-month-old males found that inulin supplementation for one month alters the microbiome distinctly from both DPR and normal protein (NP) diets (^16^). This study also demonstrated that higher inulin intake during DPR is correlated with increases in FGF21 levels and improvements in metabolic outcomes(^16^). However, whether similar mechanisms are active in both male and females during aging, or whether FGF21 is required for DPR-induced microbiome remodeling during aging, remains unknown.

These promising findings in young male mice suggest that DPR may play a critical role in regulating the gut-brain axis. Nevertheless, significant gaps remain. The effects of DPR on the gut microbiome during aging, especially in females, have not been elucidated. Furthermore, the extent to which the microbiome contributes to DPR-mediated improvements in metabolic and brain health, and whether FGF21 is necessary for these microbial changes, remains to be determined. We hypothesize that DPR-induced microbial remodeling may be a key driver of neuroprotection during aging. To address our hypothesis, we evaluated the impact of low-protein (LP) diets in aged female mice, with the objectives of characterizing microbiome adaptations to dietary protein restriction, determining their association with brain health and cognitive function, and investigating the relationship between microbiome remodeling and FGF21-mediated metabolic responses. Our study concludes that LP diets remodel the aging microbiome, induce distinct molecular and structural changes in the brain, and that these effects are correlated with the induction of FGF21. Our findings uncover a previously unrecognized axis linking dietary protein restriction, the gut microbiome, FGF21, and brain aging, with implications for sex-specific therapeutic strategies targeting healthy aging.

## Methods

### Animals and Diets

All animal procedures were approved by the USC Institutional Animal Care and Use Committee (IACUC) and were performed following the guidelines and regulations of the NIH Office of Laboratory Animal Welfare. Female C57BL/6 mice (WT, Jackson Lab) were used in all studies. Diets were formulated and produced by Research Diets as previously described(^17^) and were designed to be isocaloric by equally varying protein and carbohydrate while keeping fat constant. Normal-protein control diets (CON) contained 20% casein (by weight) as the protein source, while the low protein diet (LP) contained 5% casein. All diet compositions are provided in supplemental table 1. Aging mice that were clinically determined as being unable to thrive were removed from the study and euthanized. At the end of the study, mice were euthanized during the mid-light cycle in the fed state (unless otherwise noted) using acute exposure to CO_2_ followed by transcardial perfusion with 20 ml ice-cold 0.1M PBS, at which time intestinal tissue, liver, fat pads (inguinal, mesenteric, and gonadal), were collected. Tissues were collected and snap-frozen in liquid nitrogen for further analysis. The brains were hermisected. One hemibrain was fixed for 48 hours in 4% PFA in 4% parafmormaldehyde/0.1M PBS. After 48 hours, the brains were stored in 4°C with o.1M PBS/0.03% NaN_3_. The other hemibraine was micro-dissected and snap-frozen for downstream analysis. Whole blood was collected using a cardiac draw and centrifuged at 3400rpm, to separate plasma from white and red blood cells

### Experimental Design

To investigate whether late in intervention of LP-diet reduces the burden of gut dysbiosis and related factors on metabolic health and brain health during aging, female C57BL/6J mice were entered into the study at approximately 16 months of age and group-housed (4 per cage) at room temperature (23 °C). Mice were randomly assigned to either diet (10-14 mice/diet): normal-protein control (NP) and LP *ad libitum (supplemental table 1)*. Bodyweight and food intake were recorded weekly throughout the experiment. Body composition was measured via TD-NMR (Bruker Minispec) at the start (16 months of age) and near the end (22 months of age) of the study. Glucose homeostasis was evaluated by a glucose tolerance test at 18 months of age. After 26 weeks of dietary manipulation, mice were euthanized at 22 months of age for blood and tissue collection for further processing and analysis. The experimental timeline is presented in Figure 1A (created by BioRender).

**Figure 1.**
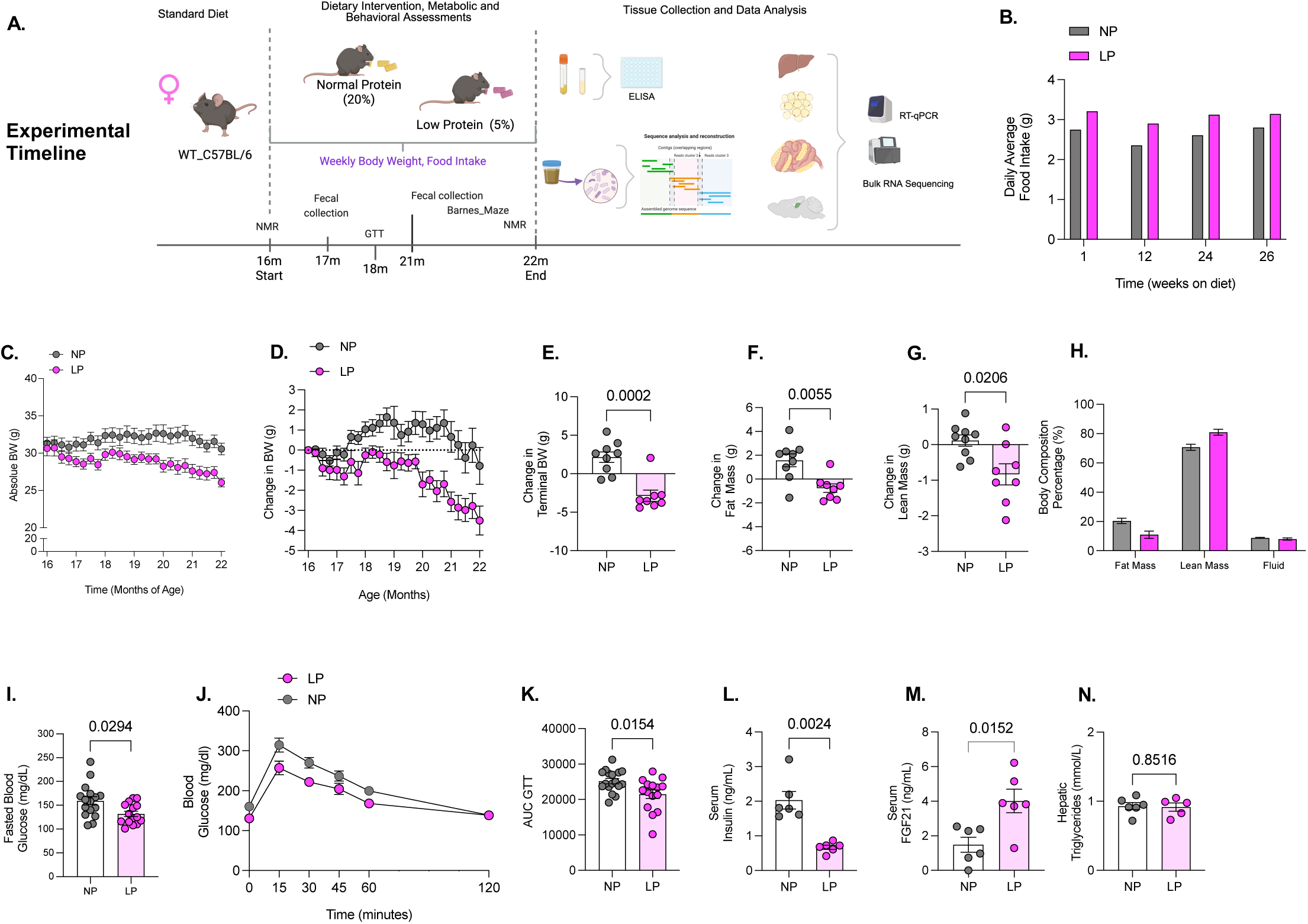
Low-protein diet late in life prevents age-related metabolic decline in females. **A** Graphical methodology: C57BL/6 (WT) female mice were placed on NP or LP at 16 months of age various metabolic endpoints on BW, body composition, glucose homeostasis throughout the feeding phase of the study as indicated (*n*=14 NP and *n*=12 LP mice /group), fecal collection at 1 month and 6 months after start of diet (*n*=3 mice per diet), Barnes maze (*n*=14 NP and *n*=12 LP mice /group), followed by tissue collection at 22m of age. **B)** Average Food Intake**. C)** Absolute weight gain over time from initiation of diet. **D)** Change in body weight over time. **E)** Terminal change in body weight. **F**) Fat gain at 21 months of age. **G)** Lean gain at 21 months of age. **I** Fasting blood glucose at 18months of age. **J** Glucose tolerance test conducted at 18 months of age (*n*=12 mice/diet). **K)** Area under the curve glucose for the GTT. **L)** Serum insulin levels at 22 months of age. **M** Serum FGF21 levels at 22 months of age. **N)** Hepatic triglyceride levels at 22 months of age (*n*=5 mice per diet). Statistical analyses were conducted using All values are mean ± SEM, with significant main effects of protein by unpaired *t*-test

### Glucose Tolerance Test (GTT)

Sixteen-hour-fasted mice underwent GTT by i.p. injection with 2g glucose per kg of BW. Blood glucose levels were measured at 0, 15, 30, 45, 60, and 120 min via a handheld glucometer (Accu Check; Roche Diabetes Care, Inc. Indianapolis IN). The data for GTT are represented as mg/dL and as the area under the curve (AUC). Fasted glucose levels are reported at the time of GTT.

### Immunoassay determination of FGF21 and Adiponectin

Plasma concentrations of FGF21 (no. RD291108200R, Mouse and Rat FGF-21 ELISA, BioVendor) and insulin (Catalog #80569, Mouse Crystel Chem) were determined via ELISA according to the manufacturer’s recommended protocol.

### Determination of hepatic triglycerides

Concentrations of hepatic triglycerides were measured in hepatic homogenates with a colorimetric assay (no. 10010303, Cayman Chemical) according to the protocol recommended by the manufacturer.

### Real-time PCR

RNA extraction and real-time PCR were conducted as previously described (^17^). Liver, depots of subcutaneous adipose-(inguinal fat - iWAT) and visceral adipose-tissue (gonadal-gWAT, and mesenteric - mWAT) of C57BL/6 female mice fed either normal protein-(NP) or low protein-(LP) diet (*n* = 5-12 per group) were used for total RNA extraction using TRIzol reagent following the manufacturer’s protocol (Invitrogen), with the addition of a RNeasy Lipid Tissue Mini Kit (QIAGEN) for adipose tissue depots. RNA purity and quantity were determined by spectrophotometry using a Nanodrop One Spectrophotometer (Thermo Scientific™ Invitrogen. cDNA synthesis was performed with iScript (BioRad) and mRNA was quantified on the ABI 7900 platform using the ABI SYBR Green PCR Master Mix in optical 384-well plates (Applied Biosystems). Primer pairs were designed using the IDT RealTime qPCR Primer Design tool to span an intro-exon boundary (supplemental table 2). Target gene expression was normalized with beta actin for liver cyclophilin for adipose tissue as the endogenous control.

### Transcriptomic and Bioinformatics Analysis

Total RNA concentrations were diluted to 50 ng/µl and submitted to Novogene for high-throughput RNA sequencing. Stranded libraries were generated using ribo-depleted total RNA with a minimum RNA integrity number (RIN) score ≥ 9. Libraries were sequenced with paired-end 150 bp reads, with three biological replicates per diet group. After initial quality control by Novogene, paired-end sequencing on Illumina platforms resulted in the generation of raw FASTQ files. These files were assessed for read quality, adapter contamination, and sequence duplication. Adapter sequences and low-quality bases were trimmed using Trimmomatic (v0.39) with default parameters. The cleaned reads were aligned to a reference mouse genome (GRCm39) using the splice-aware aligner STAR (v2.7.9a). Mapping rates and coverage uniformity of the aligned reads were assessed with Samtools (v1.15) and Qualimap (v2.2.2). Raw gene count quantification was performed using featureCounts (Subread package v2.0.3) against the GENCODE mouse gene annotation (release M30).

Our study includes dot plots of Gene set enrichment analyses (GSEA) to demonstrate diet effects. During the preprocessing stage, genes with low counts were filtered out using the DESeq2 R package (v1.38.1) automatic filtering process. Differentially expressed genes were identified using DESeq2 with a false discovery rate (FDR) cutoff of adjusted p-value (p-adj) < 0.05. Additionally, genes with log₂ fold changes of ≤ −0.5 or ≥ 0.5 were considered significant based on DESeq2 analysis results, with this threshold applied as part of the analysis workflow. An absolute threshold of 0.5 was selected for hippocampal and hypothalamic sequencing data in accordance with literature standards. Prior work has demonstrated that gene expression changes in the brain are typically more subtle than twofold (log₂ fold change = 1). Principal component analyses were performed to visualize differences across samples. Functional Gene Ontology (GO; Biological Process) enrichment analyses were conducted using the clusterProfiler package (v4.6.0) to identify pathways enriched for diet–gene interactions.

### Shotgun Sequencing and Microbiome Analysis

Fecal samples were collected mid-morning during routine handling after one month of diet (17 months of age) and after 5.5 months of diet (21.5 months of age), two weeks before tissue collection. Two fecal pellets from each animal were placed in Stool Nucleic Acid Collection and Preservation System tubes (Norgen Biotek, Catalog No. 63700) and shipped to Transnetyx (Cordova, TN, USA) for analysis. DNA extraction was performed using the DNeasy 96 PowerSoil Pro QIAcube HT extraction kit (Qiagen, Catalog No. 47021) which was optimized by Transnetyx for reproducible extraction of high-quality, inhibitor-free genomic DNA. Library preparation was performed using the KAPA HyperPlus library preparation protocol with unique dual-indexed (UDI) adapters to prevent read misassignment. Libraries were pooled and sequenced on an Illumina NextSeq 2000 platform (2X150 bp paired-end reads, 2 million read pairs per sample). Raw FASTQ data were analyzed using the One Codex database containing >148,000 microbial reference genomes. Taxonomic classification was performed using k-mer-based analysis (k=31) with automatic artifact filtering to remove sequencing artifacts and contamination while retaining low-abundance reads. Species-level abundance estimation was calculated based on sequencing depth and coverage across reference genomes.

### Behavior

The Barnes Maze test was conducted to assess learning and memory at 25 weeks of diet, one week prior to the end of the study. Details of the modified protocol used during this study are as those previously published(^18, 19^). The maze consists of 20 evenly spaced holes (5 cm diameter) along the circumference of a circular platform. Following one day of habituation under red light, mice underwent four days of training with three 3-minute trials per day, with 15 minutes between trials. Before each testing day, mice were habituated in the room for 30 minutes. At the start of each test, mice were placed at the center of the platform inside an opaque cylinder, which was lifted after 10 seconds. Mice had 3 minutes to locate the designated target escape box (11 cm L × 5 cm W × 5 cm H). During the test, a buzzer and light remained active and were turned off immediately upon the mouse entering the escape box, where they remained for one minute to reinforce learning. If a mouse failed to find the escape hole within 3 minutes, it was guided to the target box, and the buzzer and light were turned off for one minute before the animal was returned to its cage. Three decoy boxes (5 cm L × 5 cm W × 2.5 cm H) were positioned around the platform during training days. The probe trial followed the same protocol with two important modifications. First, all holes were fitted with decoy boxes, and escape latency was measured as the time required to reach the previously designated target location (indicated by a nose poke into that hole). Second, mice received only one 3-minute trial. All trials were recorded using Noldus Ethovision XT software (version 14).

### Quantification and Statistical Analysis

Data for body weight, food intake, metabolic endpoints, gene expression, and Barnes Maze were analyzed using Prism by *t*-test. All data are expressed as mean ± SEM, with a probability value of 0.05 considered statistically significant.

## Results

### Low-protein diet late in life prevents age-related metabolic decline in females

Diets low in protein have been widely shown to recruit the liver-derived hormone FGF21 to improve metabolic health and extend lifespan; however, most preclinical evidence has been generated in male mice(^17, 20–25^). In humans, reduced protein intake has also been associated with improvements in metabolic health, although in older adults, protein-restricted interventions may increase mortality risk(^9, 26–30^). Here, we assessed the interplay between dietary protein restriction (DPR) and advanced age on metabolic health outcomes and examined whether gut-health adaptations contribute to brain health during aging in female C57BL/6 mice (Figure 1A).

Consistent with previous data in male mice that initiated DPR at 16-months of age until 22-months of age (^31^), aging female mice adapt to LP-feeding with increased food intake, reduced body weight that resulted in overall reduced gain in body weight over time, and final weight (Figure 1B-E). Body composition analysis show that LP-diet reduced gains in absolute fat and lean mass (Figure 1F-G). Although LP-diet reduced both fat and lean mass, the reduction in fat mass was larger than that of lean mass, such that lean mass relative to body weight (percent body fat) was increased in LP-fed mice compared to NP-fed controls (Figure 1H).

There is substantial evidence that low-protein fed mice have enhanced glucose clearance during glucose tolerance challenge compared to normal-protein fed counterparts throughout lifespan(^31^). Here, in female mice at 18-months of age, LP-diet enhanced glucose homeostasis on measurements of reduced fasted blood glucose levels and improved glucose clearance during GTT as revealed by area under the curve (Figure 1J-K). The reduction of body fat and improved glucose homeostasis during low protein diet are reflected in the gero-protective hormones, insulin, insulin-life-growth factor, and fibroblast growth factor 21 (FGF21). Here, consistent with our previous studies in male mice, LP-fed female mice have reduced serum levels of insulin and increased FGF21 levels at 22-months of age (Figure 1L and 1M). Lastly, there was no significant difference in hepatic triglycerides by either diet in 22-month-old female mice.

### Aged females adapt to low-protein diet on tissue-specific responses to amino acid restriction and improved thermogenesis

Through nutrient sensing and endocrine signaling, the liver and adipose tissue coordinate whole-body metabolism and influence healthy aging. The liver, specifically, plays a central role in sensing amino acid availability and coordinating systemic metabolic adaptation to fluctuations in dietary protein intake. In response to reduced protein consumption, the liver adapts by inducing the expression of *Fgf21* and genes involved in amino acid biosynthesis and the amino acid response pathway. We and others have demonstrated that hepatic expression of *Fgf21* and markers of the amino acid response pathway associated with amino acid sensing and synthesis are upregulated during dietary protein restriction (DPR) in both young and aged male mice(^31–33^). Here, female mice at 22-months of age, respond to LP-diet on increases in hepatic *Fgf21* and markers of amino acid sensing, there was no difference in neither *Fgf21-receptor* or *Klb* gene expression (Figure 2A-E).

**Figure 2.**
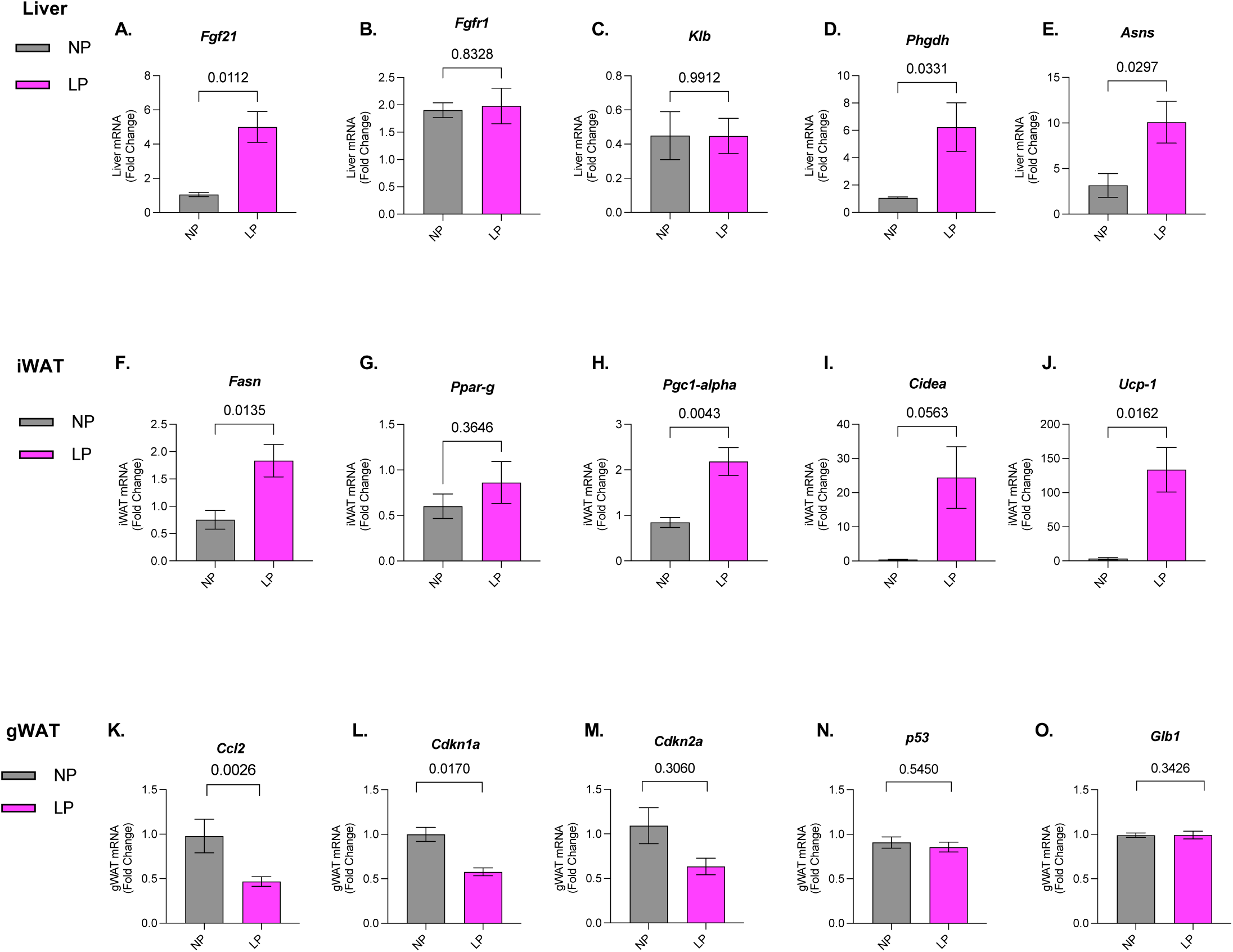
Females respond to low-protein diet on increased hepatic FGF-21 signaling and remodels white adipose tissue on improved thermoregulation and reduced SASP-related gene expression. **A-E.** Fold change of amino acid restriction markers in liver. A) *Fgf21.* B) *Fgfr1.* C) *Klb.* D) *Phdgh.* E) *Asns*. F-J. Fold change of thermogenic markers in iWAT. F) *Fsan* G) *Ppar-g* H) *Pgc-1alpha* I) *Cidea* J) *Ucp-1.* K-O) Fold change of SASP-related markers gWAT. **K.** Monocyte chemoattractant *Ccl2.* Markers of Cell cycle arrest L) *Cdkn1a* M) *Cdkn2a* and **N)** tumor protein *p53*. **O)** Senescence associated lysosomal beta-D-galactosidase *Glb1*. All values are mean ± SEM, with significant main effects of protein by unpaired *t*-test, for gene expression *n*=6 mice per diet.

Recent and new data demonstrate that the hepatic adaption to DPR on increased circulating FGF21 signals to the brain correlates with remodeling of the white adipose tissue on improved markers of thermogenesis such as *Pgc-1*, *Ucp1*, and *Cidea*. Moreover, we and others recently reported that mice deficient of FGF-21 signaling lacked adipose tissue remodeling on thermogenic reprogramming in the iWAT of 22-months of age male mice(^24, 31, 34^). Consistent with our previous data in LP-fed male mice, LP-diet, at 22 months of age in female mice increased marker of *de novo* lipogenesis *Fasn* and lipid oxidation *Pgc-1α,* although there was no difference in adipogenesis marker *Ppar-γ* (Peroxisome Proliferator-Activated Receptor Gamma) (Figure 2F-2H). Furthermore, LP-diet significantly increased makers of thermogenic programing *Cidea* and *Ucp1* in iWAT (Figure 2I and 2J), resembling reported data in male mice subjected to both short-term and long-term DPR(^8^).

Abnormal expansion of visceral white adipose tissue (vWAT) during aging contributes to adipose tissue senescence and is associated with a higher risk and incidence of metabolic disorders that may lead to cognitive decline compared with subcutaneous white adipose tissue(^35^). Likewise, adipose senescence promotes a chronic proinflammatory milieu characterized by hypoxia, lipid overflow, endoplasmic reticulum (ER) stress, all of which further exacerbate age-related disease progression(^36, 37^). We recently reported that deficiency in FGF21-signaling led to increased markers of inflammation and senescence in the epididymal WAT (eWAT) of male WT mice at 22-months of age (^31, 34^). Therefore, to identity the impact of LP-diet on markers of senescence in 22-months of age female mice, we targeted general markers in gonadal white adipose tissue (gWAT). In gWAT, LP-diet reduced *Ccl2* (also known as monocyte chemoattractant protein-1) compared with NP-fed mice (Figure 2K). Interestingly, LP-diet reduced *Cdkn1a* compared to NP-diet, yet makers of tumor suppressor *Cdkn2a* and *Trp53* and levels of expression did not reach significance (Figure 2L-2N). Lastly, Glb*1* gene expression levels were not altered by varying dietary protein content in female mice fed either diet (Figure 2O).

### A low-protein diet protects gut health through remodeling of the adipose–intestine axis

Aging is associated with changes in the gut that negatively impact the gut barrier integrity and resident microbiota. This condition, known as gut dysbiosis induces inflammation, promotes metabolic dysfunction, impairs overall health, and contributes to age-related disease, including cognitive decline. Many studies in rodent and human highlight several daily factors, including diet, exercise, biological sex, among others that alter gut health(^38–43^). In this part of the study, we targeted three features of the gastrointestinal system, thus the mesenteric adipose tissue, large intestine, and the microbiome, to encompass a multi-tissue approach on the interaction of diet and age in NP- and LP-fed female mice.

We first explored the changes in mesenteric adipose tissue. Briefly, mesenteric adipose tissue (MAT) is a specialized visceral fat depot located within the mesentery that anchors the intestines to the abdominal wall. It functions as a synonymous endocrine and immune organ that regulates energy, intestinal barrier integrity, inflammatory responses, and systemic metabolic homeostasis. Here, like the observation of reduced markers cell arrest in gWAT, LP-diet reduced *Cdkn1a* compared to NP-diet in mWAT. There was no difference in senescence related markers *Cdkn2a* and *Trp53.* In the adipose tissue, toll-like receptors (TLRs) function as a bridge between innate immune responses and metabolism that can induce chronic low-grade inflammation that is often related to insulin resistance(^44, 45^). Rodent studies show that *Tlr4* is commonly upregulated during diet induced obesity and aging, and that mice with ablation of *Tlr4* signaling during aging and obesity during aging are protected from diet-induced metabolic decline and have reduced gene expression on makers of inflammation such as *Il-6*, *Cdkn1a, and Cdkn2a*(*^46^*). While these studies highlight the role of *Tlr4* in conditions of diet and aging, these studies were conducted in male mice. Here we show that neither diet, NP nor LP, altered gene expression of *Tlr4*. Interestingly, LP-diet reduced *Il-1β*, a marker leukocyte activation in response cell damage, and trended toward lower expression of *Il-10 (*interleukin 10), a marker of suppression of immune responses during chronic inflammation to maintain systemic or tissue health(^47–49^).

Beyond macronutrient composition, habitual dietary patterns exert long-term effects on microbiome structure, while short-term dietary changes can rapidly alter microbial activity and metabolite production (^4^). Diet also interacts with other factors such as age, medication use, lifestyle, and host genetics to influence gut health outcomes(^4, 6, 50, 51^). Rodent studies show that dietary protein restriction induces metabolic adaptations through the hepatic FGF21 pathway, and emerging evidence suggests that these host responses are mediated and modulated by adaptive changes in the gut microbiome(^16^). Other studies demonstrate that adaptive responses to dietary protein restriction are mediated by the gut microbiome, which remodels host metabolism through microbial metabolites and bile acids that activate adipose FXR signaling and hepatic FGF21 production(^52^).

Consistent with these findings, finishing pigs, both barrows and gilts, fed a low-protein diet exhibited improved feed efficiency (lower feed-to-gain ratio) without compromising growth performance, suggesting that protein restriction can enhance metabolic efficiency, potentially through alterations in gut microbial composition and metabolite production(^53^). In this same study, low-protein fed gilts displayed lower circulating histidine concentrations, which correlated with reduced expression of immune and inflammatory pathways identified through KEGG analysis. While many studies establish the relationship of diet and gut flora, most of these findings are in young male mice with short-term feeding paradigms. Accordingly, given our interest in female-specific adaptations to DPR, we sought to evaluate changes in the gut microbiome following both short-term and chronic exposure to diets with varying protein content across aging. After 1-month of diet, at 17 months of age, LP-diet increased the relative abundance of *Akkermansia* by ∼18% more than NP-diet and correlated with biological processes related to the integral membrane component (Figure 3G and 3H). Interestingly, at 22 months of age, LP-diet shifted and increased the relative abundance of *Faecalibaculum* by ∼30% compared to NP-diet (Figure 3I). And that this shift in the microbiome aligned with upregulated processes related to hormone signaling, integral membrane component, and metal ion binding in LP-fed mice compared to NP-fed control counterparts (Figure 3J).

**Figure 3.**
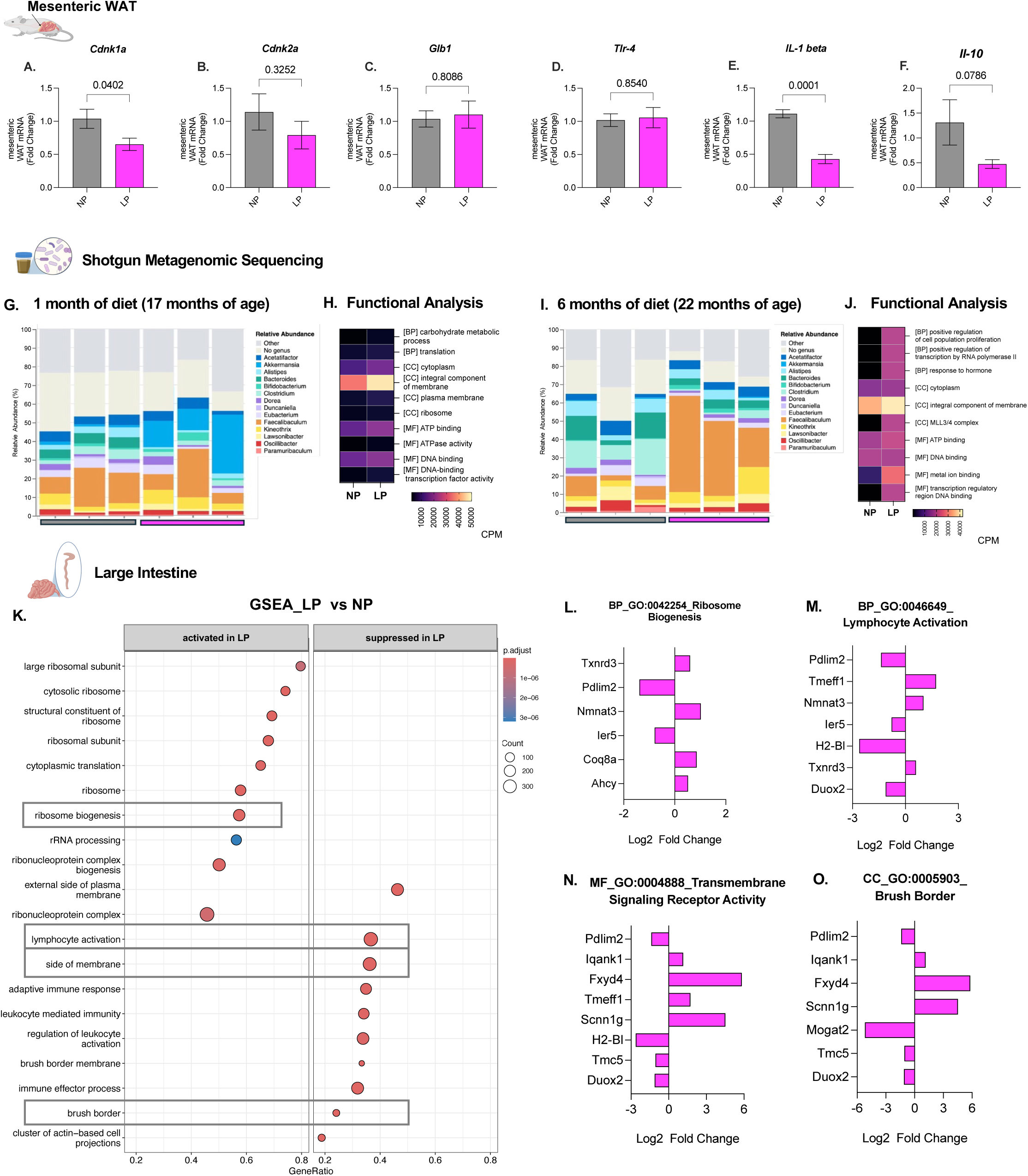
Low protein diet protects gut health on reduced age-related markers of senescence, enhanced microbiome diversity, and remodels large intestine to preserve electrolyte regulation during aging. **A-F.** Fold change of SASP-related markers mWAT, for gene expression *n*=6 mice per diet. **A.** Monocyte chemoattractant *Ccl2.* Markers of Cell cycle arrest B) *Cdkn1a.* C) *Cdkn2a.* D-E) Innate markers of inflammation. **D)** Maker for LPS detection Toll-like receptor 4 *Tlr4*. E) Interleukin1-beta, *IL-1β*. F) Interleukin 10, *IL-10.* G) Bar plot of average relative abundance at the genus taxonomic level after 1 month of diet, top 15 phyla are shown, *n*=3 NP and *n*= 3 LP. **H)** Enrichment analysis of taxonomic profiles of microbiome at 22 months of age. **I)** Bar plot of average relative abundance at the genus taxonomic level after after 6 months of diet, top 15 phyla are shown *n*=3 NP and *n*= 3 LP. **J)** Enrichment analysis of taxonomic profiles of microbiome at 22 months of age. **K)** Gene set enrichment analysis (GSEA) of large intestine at 22 months of age. **L-O)** Fold change of genes from selected GESA GO-Terms from large intestine bulk-rna-Seq in LP-fed (*n*=4) female mice at 22 months of age. All values are mean ± SEM, with significant main effects of protein by unpaired *t*-test.

Aging is associated with structural and functional changes in the large intestine, including alterations in epithelial turnover, barrier function, and immune responsiveness(^1^). Concurrently, changes in the microbiome may affect the expression and function of brush border-associated proteins, compromise epithelial integrity, and contribute to dysregulated immune responses, ultimately impacting gastrointestinal health during aging (^2, 54–56^). In addition to age-related change in the large intestine, females experience hormonal fluxes that may contribute to the changes in gut health, thus the role of estrogen-type signaling within the colon has been shown to be protective of intestinal permeability (^57–59^). Here via an unbiased approach, at 22-months of age, LP-diet activated pathways of ribosomal function and gene signatures of biogenesis that support antioxidant defense (*Txnrd3, Coq8a, Ahcy*) (Figure 3K and 3L). Interestingly, LP-diet suppressed pathways related to lymphocyte activation, transmembrane signaling, and the brush boarder that through the DEGs of the selected pathways support intestinal host-interaction defense on reduced gene signatures of DNA damage and cellular stress (*Ler5, Duox2, Tmc2,* and *Mogat2*) and increased gene signatures of epithelial function, ion transport, and mucosal homeostasis (*Tmeff1, Fxyd4, and Scnn1g*) (Figure 3K and 3M–3O).

These data indicate that the well-established benefits of DPR are effective on gut health. Compellingly, aged female mice adapt to a LP-diet on persevering gut health on reduced markers immuno-senescence, colonization of beneficial microflora, and perseveration of colon function related to electrolyte regulation and homeostasis of the brush border.

### Hippocampal adaptation to a low-protein diet is associated with improved learning behaving during aging

In the Barnes maze, LP diet improved performance, with a significant effect on the probe day, a measure of memory retention. LP animals showed marginal improvement on training days, but these differences were not statistically significant (Figure 4A-C). On the probe day, however, LP animals demonstrated faster primary escape latency and significant reductions in errors (Figures 4D-E). To understand the molecular adaptations corresponding to these memory improvements, we microdissected hippocampal tissue and analyzed RNA expression signatures. Gene set enrichment analyses revealed that LP diets suppress vascular, endocrine, and inflammatory programs including blood vessel development, vasculature development, cytokine response, glucocorticoid response, and hormone responses (Figure 4G). In contrast, LP diets activated processes relating to DNA replication and repair, microtubule motor activity, and cellular extravasation, suggesting inductions of protective mechanisms. These processes help cells respond to damage and activate repair pathways to maintain cellular integrity (Figure 4G). We further analyzed the genes driving these processes and founded enrichments in *Blm*, *Tipin*, *Cdc40*, and *Xpa* alongside suppression of *Gadd45b*, *Xbp1*, *Jade2*, *Cdkn1a*, *Tiparp*. These patterns indicate reduced stress-responsive signaling and shifts toward repair and proliferation of progenitor cells (Figures 4H and 4I). Consistent with previous literature, we found that genes related to glucose metabolism and mTOR signaling were downregulated, further supporting the evidence that LP diets more efficiently maintain glucose homeostasis and suppress mTOR-dependent cellular stress (Figure 4J).

**Figure 4.**
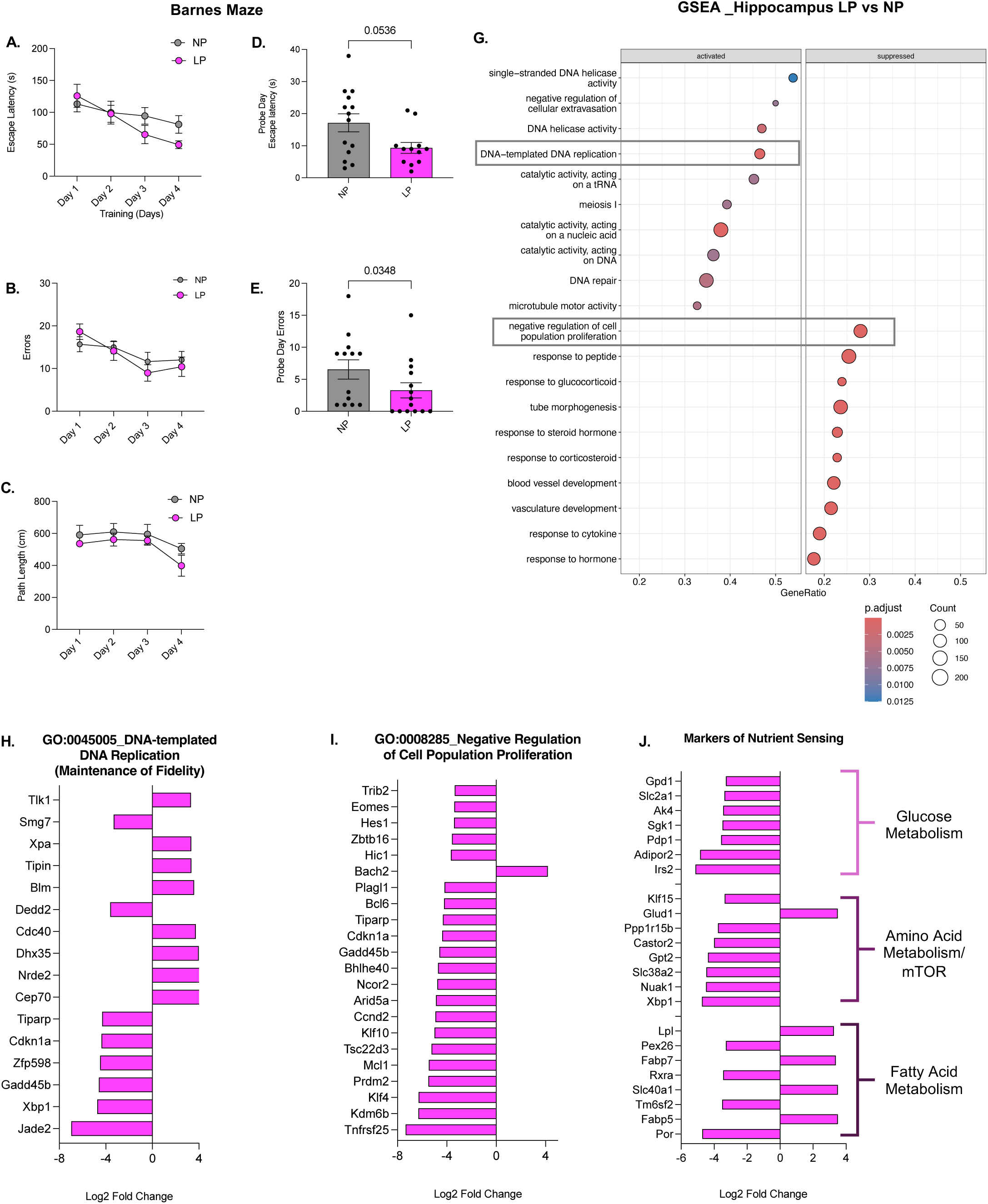
Low protein diet in aged females yield improved learning and memory is associated with improved hippocampal DNA integrity, transcriptional regulation and energy metabolism. **A-E**) Barnes maze training days, *n* = 14 normal protein (NP) diet and *n*=13 low-protein (LP) diet. **A)** Escape latency during the Barnes maze training days. **B)** Total errors during the Barnes maze training days. **C)** Path length during the Barnes maze training days. **D)** Escape latency on probe day. **E)** Escape latency on probe day. **G)**Gene Set Enrichment Analysis for GO: Biological Processes in hippocampus of NP (*n*=4) and LP (*n*=4). **H-J)** Fold change of genes from selected GESA GO-Terms from hippocampal bulk-rna-Seq in LP-fed (*n*=4) female mice at 22 months of age. Statistical analyses were conducted using a t-test. All values are mean ± SEM, with significant main effects of protein.

Finally, enrichments in *Fabp7*, *Fabp5*, *Slc40a1*, and *Lpl* alongside downregulation of *Rxra*, *Pex26*, *Tm6sf2*, and *Por* indicate that LP-fed animals reprogram lipid metabolism to preserve and enhance cellular regulation during aging (Figure 4J). Overall, these data indicate that females during aging adapt and respond to dietary protein restriction on improved metabolic health, and that the gut-brain axis enhances the remodeling of the hippocampus on DNA integrity and regulation of uncontrollable cell proliferation is associated with improvements in memory behavior.

## Discussion

Maintaining gut health through diet has been linked to a wide range of benefits, including improved metabolic function, enhanced immune regulation, and reduced risk of gastrointestinal and systemic diseases. Therefore, dietary strategies such as increased fiber intake, consumption of fermented foods, and adherence to balanced dietary patterns (e.g., Mediterranean diet, calorie restriction) are increasingly recognized as key interventions for supporting a resilient and functionally robust gut microbiome(^60^).

In male mice, both our lab and others have demonstrated that DPR confers similar metabolic and health benefits, including improved insulin sensitivity, beneficial changes in body composition, and increased expression of the liver-derived hormone, fibroblast growth factor 21 (FGF21). FGF21, which is robustly induced by DPR, is essential for many DPR downstream effects, including improved insulin sensitivity and thermogenic remodeling or “browning” of adipose depots in an uncoupling protein 1 (UCP1) dependent manner. However, these benefits have not yet been translated in female mice, highlighting an important sex-specific divergence in the physiological response to DPR. Mechanisms behind this sex difference remain unclear but are thought to involve the distinct white adipose tissue remodeling and metabolic adaptations during the reproductively active stage. Therefore, while males exhibit a consistent beneficial response to DPR, the response in females is absent, suggesting that hormonal regulation or feedback pathways require further study. However, whether these changes occur in aged female mice remains unknown.

Regarding altering protein intake, most rodent studies have demonstrated that young female mice only respond to DPR when surgical intervention of ovariectomy are performed(^61–63^). Here we show that aged female mice respond to a LP-diet on reduced gains in body weight, fat mass, preserving lean mass per gram of body weight, awhile increases in food intake. Additional enhance measures of health are marked by improved glucose clearance during GTT, lower insulin levels, and increased circulating FGF21. We further show that hepatic adaption to LP-diet is consistent on increases in gene expression of FGF21 and markers of amino acid biosynthesis *Phgdh* and *Asns* as observed in ours and other studies (^17, 22, 24, 31, 61, 64, 65^).

The gut microbiome is an intricate ecosystem of bacteria, archaea, viruses, and fungi residing in the gastrointestinal tract that plays a fundamental role in maintaining host health throughout life(^41, 66, 67^). It contributes to nutrient metabolism, immune regulation, intestinal barrier integrity, and signaling between the gut and distant organs such as the brain and liver (^43, 68, 69^). However, biological aging is accompanied by profound shifts in microbiome composition and function, often characterized by reduced microbial diversity, loss of beneficial taxa (such as *Faecalibacterium* and *Akkermansia*), and an increase in pro-inflammatory or pathogenic species(3, 6). Here we investigated the gastrointestinal system with a targeted focus on mesenteric adipose tissue, large intestine, and the microbiome, to comprehend the gut milieu in response to varying dietary content during aging we targeted three features of the gastrointestinal system, thus the mesenteric adipose tissue, large intestine, and the microbiome, to establish the impact of diet and age in NP-and LP-fed female mice. In female mice at 22-months of age, LP-diet reduced senescence-related makers in the mesenteric adipose tissue compared to NP-fed controls. Further investigation of the gut established that after 6-months of LP-diet increased abundance of *Faecalibacterium* and this shift in the flora modified pathways that reflect response to hormone, intestinal epithelium, and ion binding. These enriched pathways compellingly suggest that LP diets induce a more metabolically active gut environment. Lastly, we leveraged bulk-RNA-seq to confirmed if our changes in the microbiome also reflected colon health on function and regulation. Our GSEA analysis show that LP-diet activated ribosome biogenesis and related gene that support antioxidant defense, as well as suppressed pathways related to immune function and transmembrane signaling that supported improved control of cell proliferation and cellular stress in aged female mice.

Gut dysbiosis, characterized by imbalanced microbial communities and increased gut permeability, is associated with neuroinflammation and protein aggregation, both hallmarks of neurodegenerative diseases such as Alzheimer’s and Parkinson’s (^68, 70^). Moreover, while diet has long been recognized as a modifiable determinant of the gut-brain axis during aging, the role of DPR in shaping the gut-brain axis during aging females remains to be fully explored during normal aging conditions. Here we show that LP-diet improved memory retention in aged female mice, as evidenced by enhanced Barnes Maze performance during the probe trial. These cognitive benefits were accompanied by hippocampal transcriptional remodeling characterized by suppression of inflammatory, vascular, stress-response, glucose metabolism, and mTOR signaling pathways, together with activation of DNA repair, genome maintenance, and cellular protective mechanisms. Together, these findings suggest that dietary protein restriction promotes cognitive resilience during aging by supporting hippocampal integrity through enhanced cellular maintenance and stress resistance.

Collectively, our findings demonstrate that dietary protein restriction promotes broad physiological adaptations in aged female mice, extending beyond metabolic regulation to encompass improvements in gut, adipose, and brain health. Unlike previous studies in young females, aged females exhibited hallmark responses to low-protein feeding, including elevated FGF21, improved glucose homeostasis, reduced adiposity, and preservation of lean mass. These systemic adaptations were accompanied by reduced markers of senescence in mesenteric adipose tissue, remodeling of the gut microbiome toward taxa and functional pathways associated with intestinal homeostasis, and transcriptional signatures in the colon indicative of enhanced cellular maintenance and reduced immune activation. Furthermore, low-protein feeding improved memory retention and induced hippocampal gene programs associated with DNA repair, stress resistance, and suppression of inflammatory and mTOR-related signaling. Together, these data support a model in which dietary protein restriction promotes healthy aging in females through coordinated adaptations across the gut–adipose–brain axis, highlighting the gut microbiome and FGF21 signaling as potential mediators of age-dependent resilience and metabolic health.

